# 3D electronic implants in subretinal space: long-term follow-up in rodents

**DOI:** 10.1101/2023.07.25.550561

**Authors:** Mohajeet Bhuckory, Bing-Yi Wang, Zhijie Charles Chen, Andrew Shin, Davis Pham-Howard, Sarthak Shah, Nicharee Monkongpitukkul, Ludwig Galambos, Theodore Kamins, Keith Mathieson, Daniel Palanker

## Abstract

Photovoltaic subretinal prosthesis (PRIMA) enables restoration of sight via electrical stimulation of the interneurons in degenerated retina, with resolution limited by the 100 μm pixel size. Since decreasing the pixel size below 75 μm in the current bipolar geometry is impossible, we explore the possibility of using smaller pixels based on a novel 3-dimensional honeycomb-shaped design. We assessed the long-term biocompatibility and stability of these arrays in rats by investigating the anatomical integration of the retina with flat and 3D implants and response to electrical stimulation over lifetime – up to 9 months post-implantation in aged rats. With both flat and 3D implants, VEP amplitude decreased after the day of implantation by more than 3-fold, and gradually recovered over about 3 months. With 25 μm high honeycomb walls, the majority of bipolar cells migrate into the wells, while amacrine and ganglion cells remain above the cavities, which is essential for selective network-mediated stimulation of the second-order neurons. Retinal thickness and full-field stimulation threshold with 40 μm-wide honeycomb pixels were comparable to those with planar devices – 0.05 mW/mm^2^ with 10ms pulses. However, fewer cells from the inner nuclear layer migrated into the 20 μm-wide wells, and stimulation threshold increased over 5 months, before stabilizing at about 0.08 mW/mm^2^. Such threshold is significantly lower than 1.8 mW/mm^2^ with a previous design of flat bipolar pixels, confirming the promise of the 3D honeycomb-based approach to high resolution subretinal prosthesis.

## Introduction

Retinal degenerative diseases, such as age-related macular degeneration (AMD) and retinitis pigmentosa (RP), are among the primary causes of untreatable visual impairment. Patients with atrophic AMD lose central high-acuity vision due to the gradual demise of photoreceptors in the macula. This advanced form of AMD, called geographic atrophy, affects millions of patients: approximately 3% of the population over the age of 75, and 25%, over the age of 90^1,2^. As life expectancy extends, the number of patients that will suffer from severe vision loss (beyond the legal blindness limit of 20/200) is expected to increase. The hereditary RP originates from genetic disorders, which typically lead to irreversible loss of photoreceptors much earlier in life, eventually resulting in profound blindness at young adulthood^3^. In both conditions, however, the inner retinal neurons are preserved to a large extent, albeit with some rewiring^4,5^, which enables restoration of sight via electrical stimulation of these downstream neurons.

Several designs of retinal prosthetics have been tested in RP patients. The Argus II epiretinal prosthesis (Second Sight Medical Products, Inc, Sylmar, CA, USA) has been implanted in more than 200 RP patients for 1-3 years, and the subjects performed better in functional visual tests with the system ON than OFF, although a third of the subjects experienced serious adverse events (SAEs)^6^. None of the RP patients with Argus II gained vision to the extent that would allow ambulation without a white cane or a guide dog. In clinical trials of the Alpha IMS/AMS subretinal prostheses (Retina Implant AG, Reutlingen, Germany), out of 44 RP patients in total, nearly all reported implant-mediated light perception, but only 6 demonstrated prosthetic letter acuity, ranging from 20/2000 to 20/550 (2 patients)^7^. Further improvement in prosthetic vision is required to provide substantial benefits to patients blinded by retinal degeneration.

For atrophic AMD, the subretinal photovoltaic arrays PRIMA (Pixium Vision, Paris, France) have demonstrated both safety and efficacy in the initial clinical trial, achieving a prosthetic letter acuity in the range of 20/438 - 20/550, closely matching the sampling limit with 100 μm pixels: 1.17±0.13 pixels^8,9^. Moreover, patients reported simultaneous perception of the central prosthetic vision and remaining natural peripheral vision^7^/25/2023 10:34:00 PM. This encouraging result paves the way for developing devices with higher resolution. For a wider adoption of this technology clinically, prosthetic visual acuity should significantly exceed the remaining eccentric vision, typically no worse than 20/400 in AMD patients.

Improvement in prosthetic resolution requires efficient stimulation of retinal neurons by smaller pixels. Scaling down the bipolar pixels in subretinal planar implants like PRIMA is fundamentally limited by the electric field penetration depth, which scales with the pixel width due to the local circumferential return electrode in each pixel^10,11^. Consequently, if continuing to use this pixel design, smaller pixels will require much stronger stimuli, exceeding the safety limit of charge injection with pixels smaller than 75 μm in human retina^11^.

One promising approach to overcome such limitation is to create 3-dimensional (3D) honeycomb-like interface^11^ by elevating the return electrodes and allowing the bipolar cells to migrate into these cavities. Vertically aligned electric field polarizes the bipolar cells more efficiently due to the vertical orientation of their axons in the retina. Vertical confinement of electric field near the bipolar cells is also important for selective network-mediated retinal activation. Preliminary results have shown that retinal second-order neurons migrate into the honeycomb wells within a couple of weeks post implantation^10,12,13^. However, given the chronic nature of retinal implants, further studies are needed to investigate the long-term safety and stability of such devices in the subretinal space. These could also provide valuable insights to other designs of 3D retinal prostheses^14–18^, and potentially other neural interfaces.

Here, we present a comprehensive longitudinal study of the 3D honeycomb-shaped implants in the subretinal space in rats. We examine both, their structural integration by studying retinal anatomy, and neural excitability by electrophysiological measurements over lifetime of the animals - 32 weeks post implantation. We compare the retinal thickness, cellular migration, as well as electrical excitability across three categories of implants: honeycombs on 40 μm pixels, honeycombs on 20 μm pixels, and planar devices, illustrated in Figure 1.

**Figure 1.**
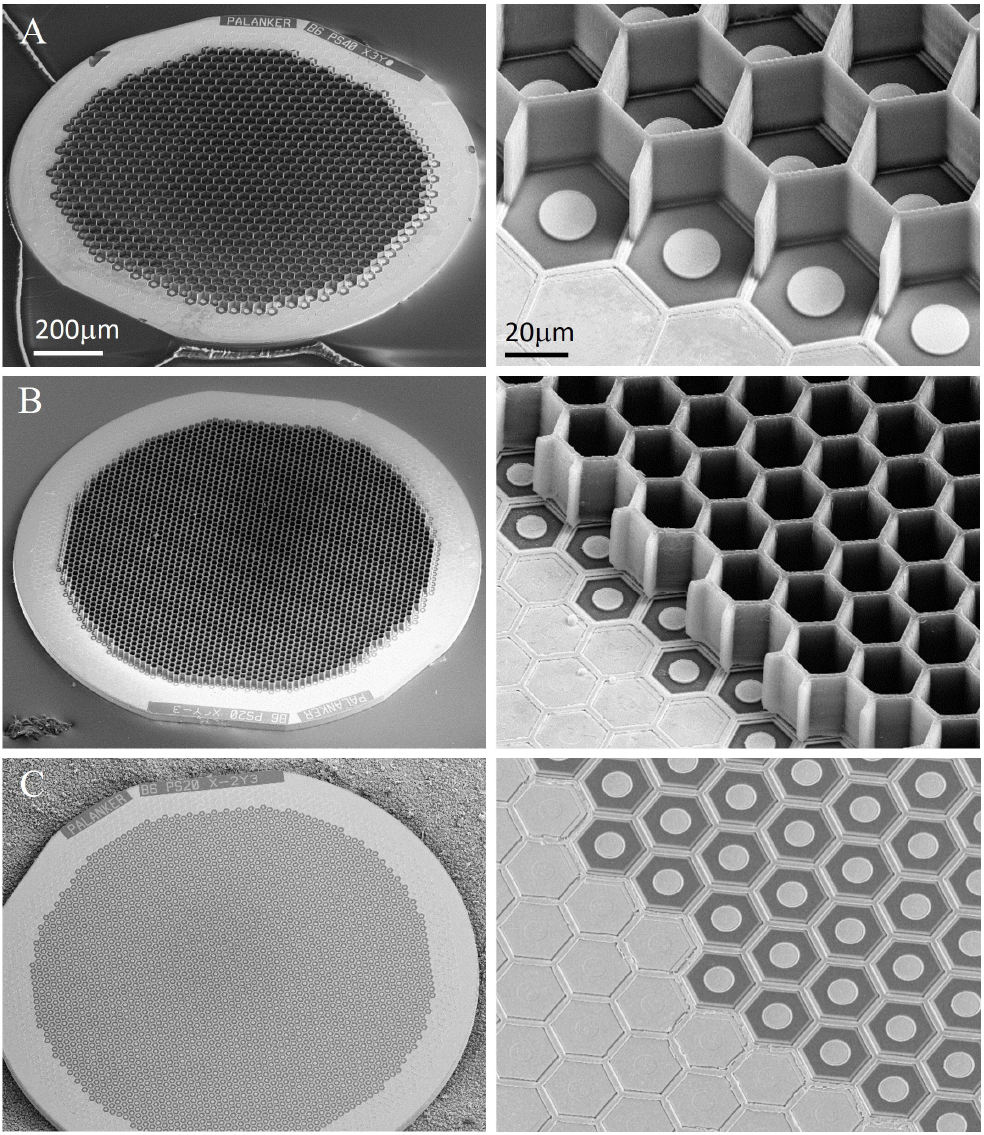
Scanning electron microscopy (SEM) of the implants. Honeycomb-shaped walls of 25 μm in height were polymerized onto planar photovoltaic arrays; (A) 40 μm pixels, (B) 20 μm pixels, (C) planar monopolar implant with 20 μm pixels.

## Results

### Retinal thickness

To assess the long-term effect of our subretinal implants (planar and 3-D) on retinal thickness, 1.5mm devices (Fig. 1 A-C) were implanted in 6-9 months old Royal College of Surgeon (RCS) rats - a model of photoreceptor degeneration. The retina and implant were monitored *in vivo* for over 32 weeks using optical coherence tomography (OCT) imaging. A hyperreflective layer, not present in the non-implanted control (Fig. 2A), becomes visible above the planar and honeycomb implants on the day of implantation (Fig. 2 E, I). This debris layer may represent the remnants of the outer plexiform layer (OPL), which becomes visible after retinal detachment and almost completely disappears after 1 week (Fig. 2F). At that time, the top of the honeycomb walls becomes visible in OCT (yellow arrow in Fig. 2J,K,L), allowing monitoring the INL migration into the wells.

Retinal morphology above all implants remained comparable to their age-matched un-implanted control. Retinal thickness, quantified using the Heidelberg software (Fig. 3A), decreased at similar rate with all implants and in the non-implanted controls (n = 3-6, two-sample t-tests, *p* = .25), which suggests an age-related mechanism of thinning in this animal model.

**Figure 2.**
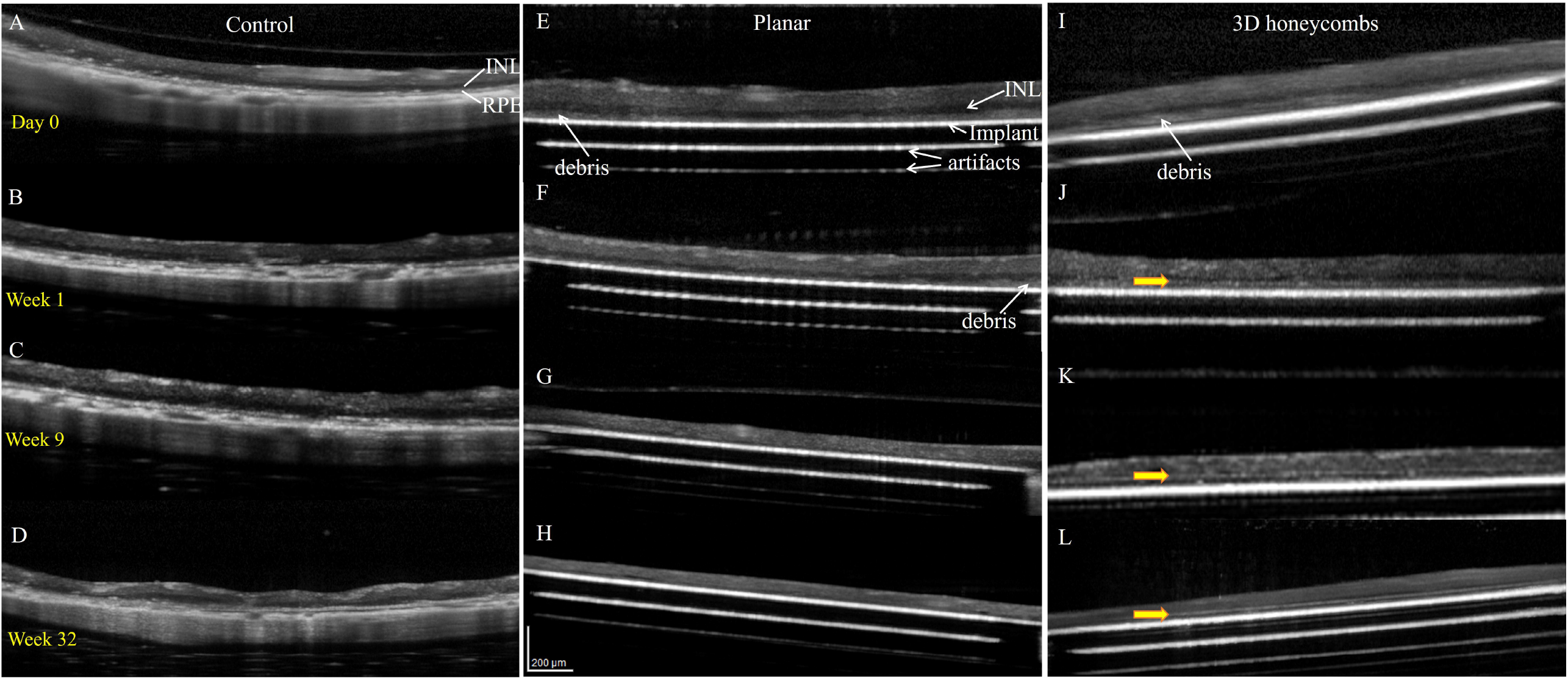
Optical coherence tomography (OCT) of the RCS rat retina at various time points: Day 0, weeks 1, 9 and 32. (A-D) Non-implanted control eyes. (E-H) Planar implants. (I-L) 3D honeycomb implants. Yellow arrows indicate the top of the honeycomb walls visible on OCT.

**Figure 3.**
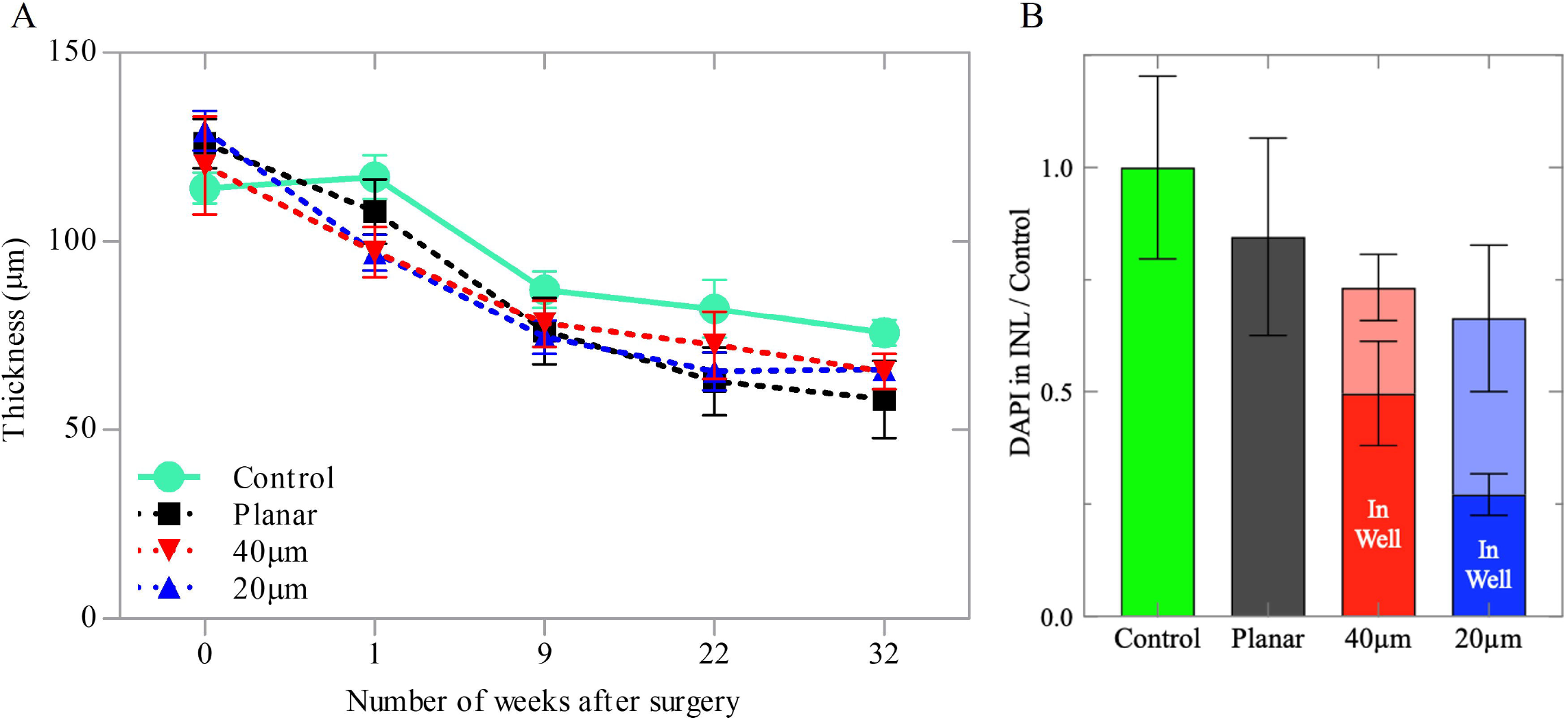
Quantification of the retinal anatomy. (A) Retinal thickness above the planar and 3D implants for up to 32 weeks post-op, compared to the age-matched non-implanted controls, measured with OCT. (B) Quantification of the cellular migration into the wells, relative to the INL in non-implanted controls, assessed with DAPI staining.

For immunohistochemical imaging, retinas from the implanted rats and their age-matched controls were collected 9 months post-implantation. The samples were labeled with DAPI and nuclei above each implant were quantified relative to the INL of non-implanted controls (Fig. 3B). Above the implants, the INL contained fewer nuclei: 0.85 ± 0.22 for planar, 0.73 ± 0.17 for 40 μm honeycomb and 0.66 ± 0.15 (two-sample t-tests, *p* = .037) for 20μm honeycomb, relative to the age-matched control. However, the differences between the planar and 3-D implants were not significant, suggesting comparable long-term compatibility of such structures in the subretinal space.

### Retinal structure

Functional success of the honeycomb implants depends on migration and long-term stability of the bipolar cells inside the wells. Morphology and location of cells was assessed by confocal imaging of the whole-mount RCS retina immuno-labelled with PKCα (rod bipolar cells) and secretagogin (cone bipolar cells) after 9 months of implantation. The retinal structure with planar implant (Fig. 4B) was comparable to that of the non-implanted control (Fig. 4A). The majority of bipolar cell somas can be seen inside the 40μm wells, but much fewer of them migrate into the 20μm wells, as quantified in Fig. 3B. Comparable total number of the INL cells (DAPI staining) in these two groups indicates that the difference is in the extent of cell migration rather than in cell death.

**Figure 4.**
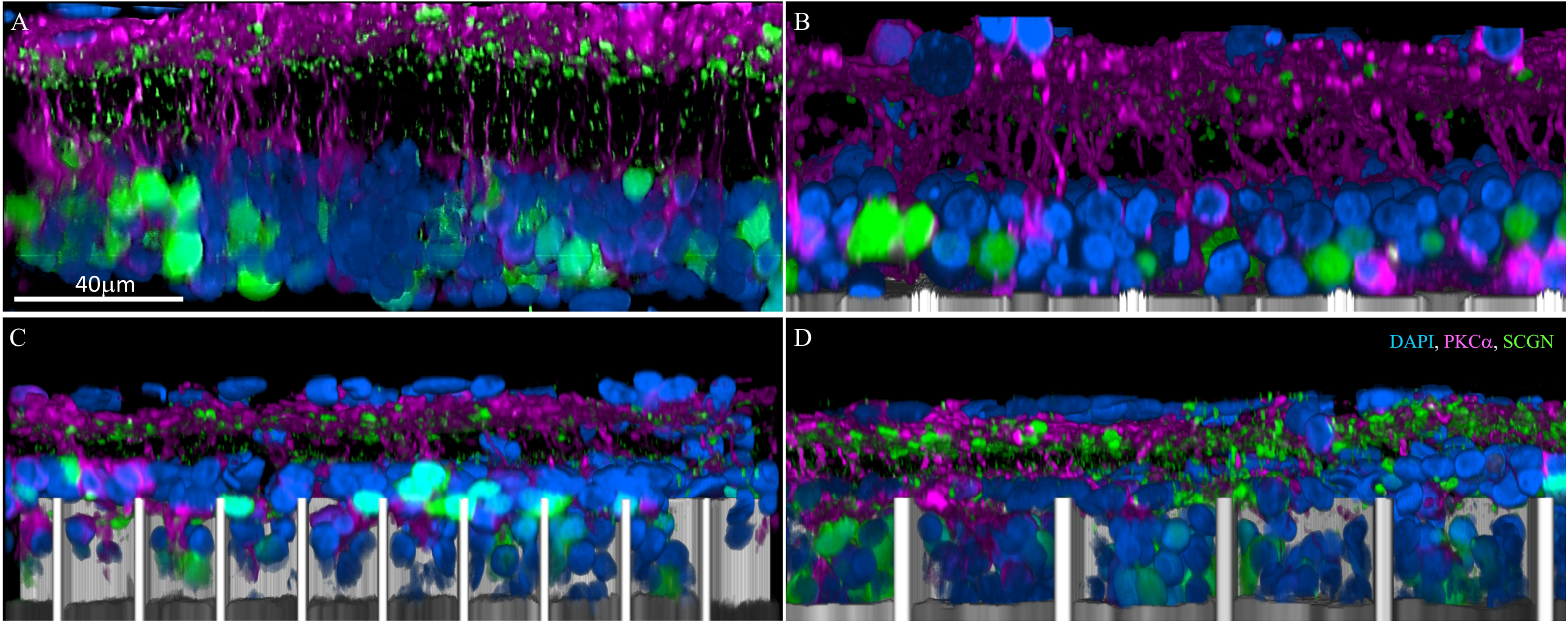
Confocal images of immuno-labelled rod bipolar (PKCα: magenta) and cone bipolar (secretagogin; green) cells. (A) Age-matched non-implanted RCS retina; (B) retina with planar implant; (C) retina above and inside the honeycomb wells of 20 μm and (D) 40 μm pixels after 32 weeks *in-vivo*. DAPI labeled nuclei are blue and the implant is grey. Scale bar = 40 μm

Another important functional requirement for the honeycomb design is discrimination between second and third-order neurons: bipolar cells should reside inside the wells to be efficiently stimulated, while the amacrine cells should remain above the return electrode on top of the walls, and thus not be exposed to strong electric field. Amacrine cell location was assessed using anti-choline acetyltransferase antibody (CHAT). As can be seen in Fig. 5C, D, amacrine cells remain outside the honeycomb wells despite the retinal thinning during the 6-9 months post-op. Furthermore, the amacrine cells morphology and the inner plexiform layer (IPL) stratification remained comparable to that with planar implants and non-implanted controls.

**Figure 5.**
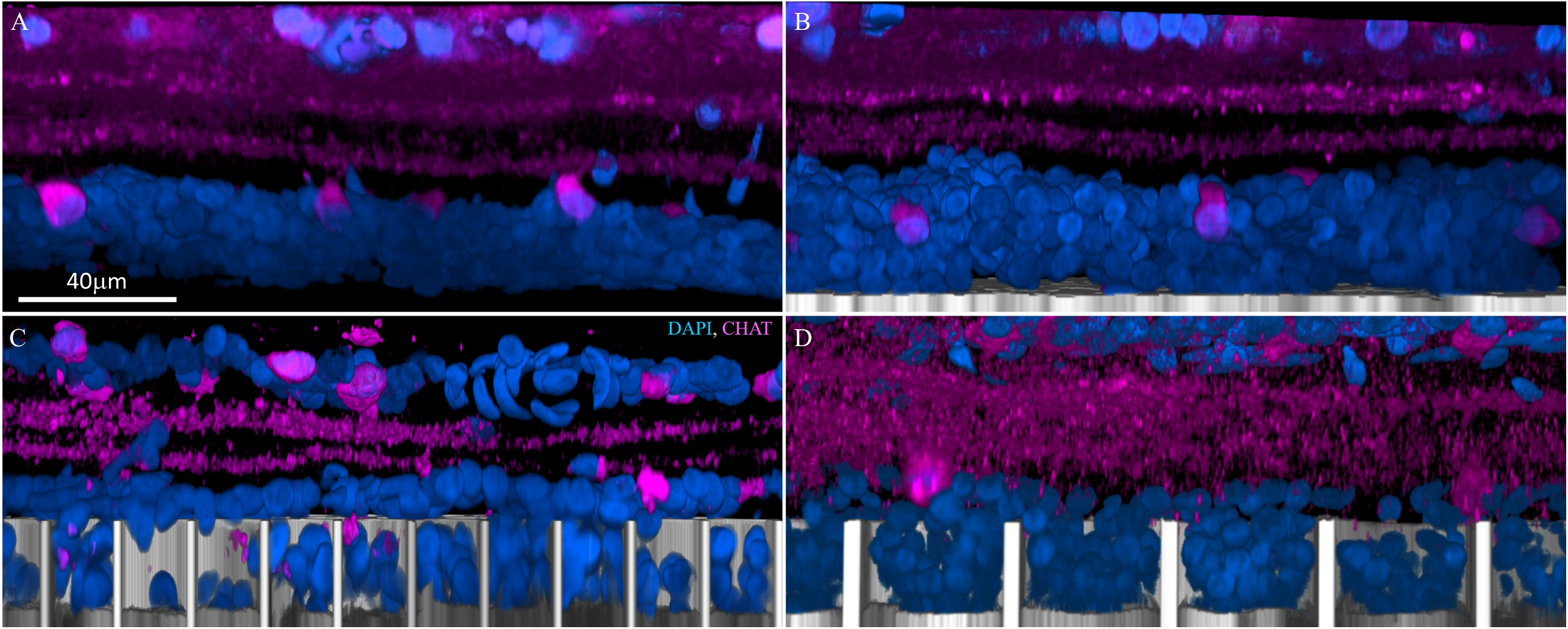
Amacrine cells immuno-labelled with choline acetyltransferase (CHAT: magenta). (A) Age-matched non-implanted RCS retina; (B) retina with planar implant; (C) retina above and inside the honeycomb wells of 20 μm and (D) 40 μm pixels after 32 weeks *in-vivo*. Amacrine cells remain above the walls for both honeycomb sizes, they retain their IPL stratification and cell body position in the INL, similar to those in the control. DAPI labeled nuclei are blue and the implant is grey. Scale bar = 40 μm.

Long-term Müller glial activation due to subretinal implants was assessed by glial fibrillary acidic protein (GFAP) staining. Comparable GFAP levels were observed between the planar, 20 and 40 μm honeycomb implants (Fig. 6 B, C, D), but all of them higher than the non-implanted control (Fig. 6A). Furthermore, a GFAP positive layer can be seen above the wells of both 20 and 40 μm pixels (Fig. 6 C, D yellow arrows), suggesting that a glial seal forms in the space left by the INL cells that migrated into the wells. This seal might be the reason why amacrine cells do not migrate into the wells (Fig. 5 C, D) even after the retina is becomes thinner.

**Figure 6.**
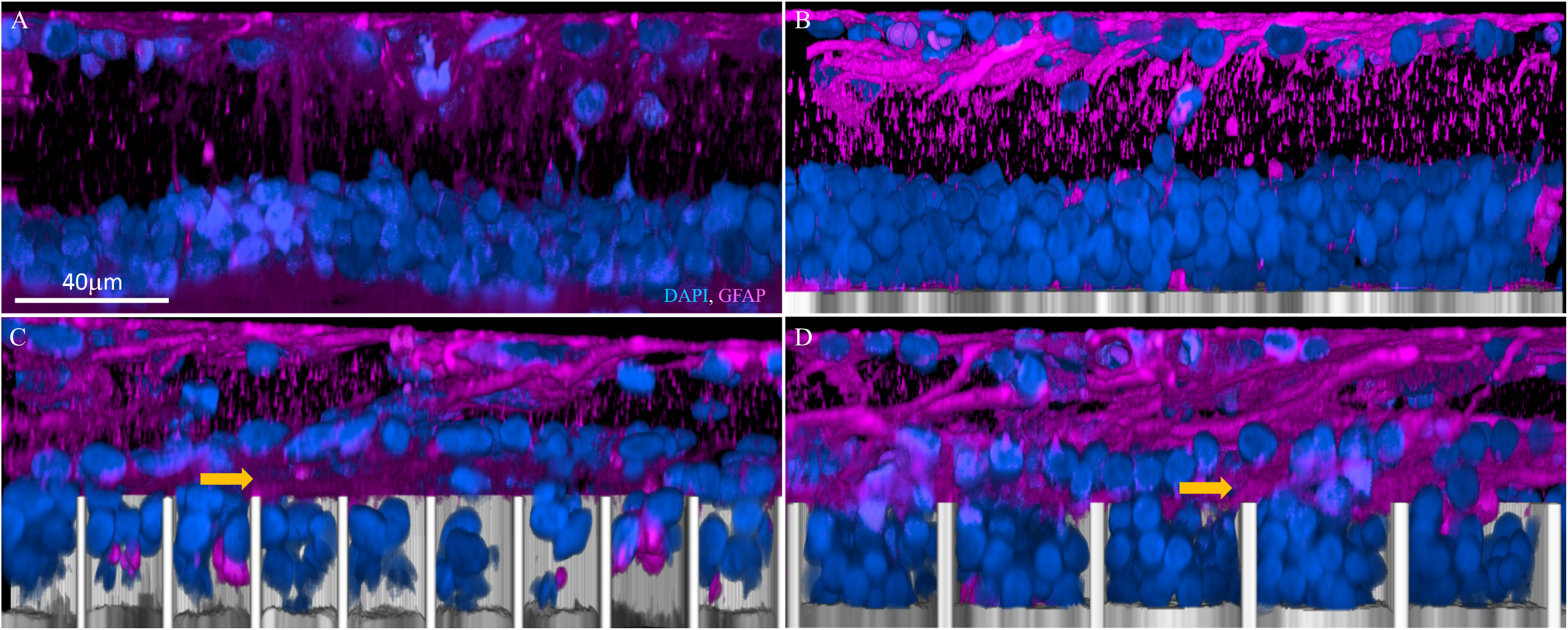
Muller cell activation marker GFAP (magenta). (A) Age-matched non-implanted RCS retina; (B) retina with planar implant; (C) retina above and inside the honeycomb wells of 20 μm and (D) 40 μm pixels after 32 weeks *in-vivo*. GFAP clusters (yellow arrow) between the cells migrated into the honeycombs and the rest of the INL in both 20 and 40 μm. DAPI labeled nuclei are blue and the implant is grey. Scale bar = 40 μm.

### Neural excitability and prosthetic vision

To assess potential changes in retinal excitability over time, we measured the visually evoked potential (VEP) in response to full-field stimulation. As shown in Figures 7 and 8, VEP amplitude decreased right after the implantation and gradually recovered over 3-4 months post-op. However, stimulation threshold remained largely constant in all three groups of implants: 3D with 40 μm and 20 μm pixels, and the flat devices (Figure 8A). Statistical tests (two-sample t-tests) among the three implant groups revealed no significant difference (p>0.05 at all the time points) between the stimulation thresholds with planar implants and 40 μm honeycombs, indicating that retinal integration with these honeycomb structures did not decrease its neural excitability. However, with 20μm honeycomb pixels, stimulation thresholds were significantly higher than the other two groups after 18^th^ week (0.01<p<0.05). After 4 months, the peak amplitude of the VEP (at 2.4 mW/mm^2^ irradiance) largely stabilized, and it exceeded the value measured at the day of implantation (Figure 8B).

**Figure 7.**
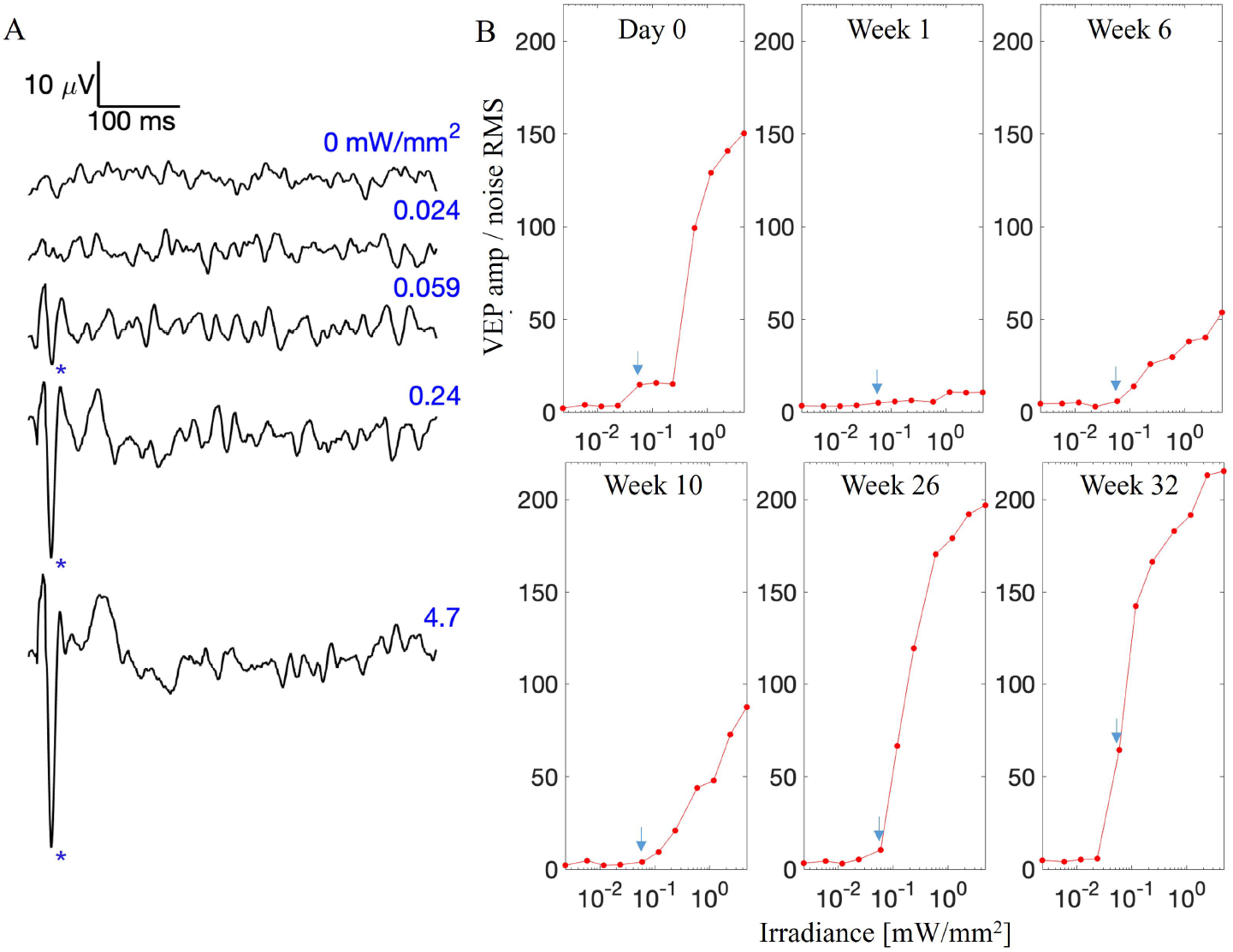
Cortical responses to subretinal stimulation over time. A) Example visually evoked potential (VEP) waveforms in response to 10ms pulses at various irradiance. Above the stimulation threshold, the characteristic N1 peak (marked by asterisks) appears in the waveform. B) VEP amplitudes as a function of irradiance, normalized by noise at selected time points. Stimulation thresholds are pointed by blue arrows.

**Figure 8.**
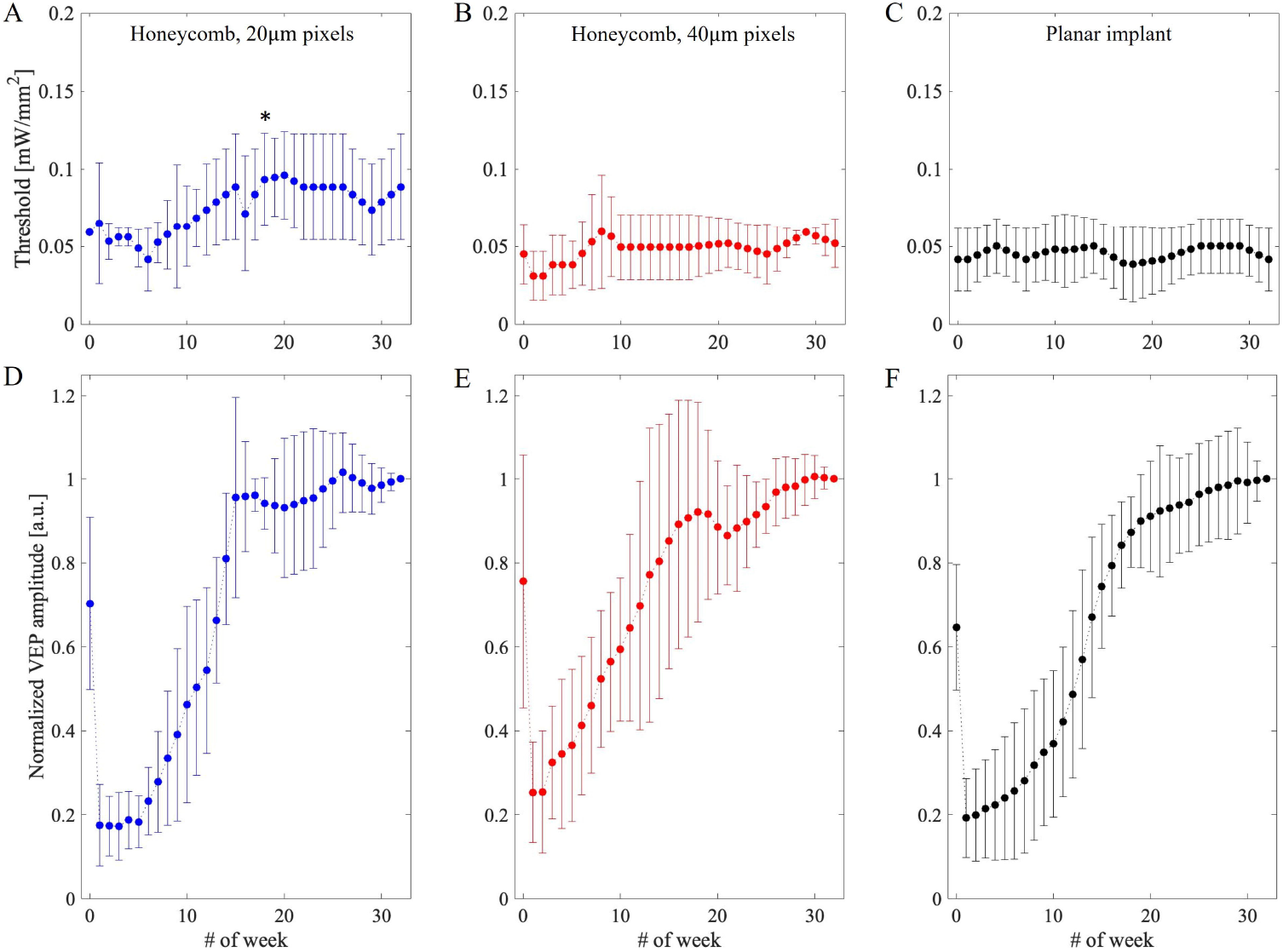
The stimulation thresholds over time - number of weeks post implantation for A) honeycomb devices with 20μm pixels, B) 3D honeycomb devices with 40μm pixels, and C) planar devices as a control. (D-F) VEP amplitudes at 2.4 mW/mm^2^ irradiance measured from the day of implantation for up to 32 weeks for 20-, 40-μm honeycomb and planar devices, respectively. Due to variability between animals, values are normalized prior to averaging by the last measurement.

## Discussion

Clinical trials with our planar bipolar pixels (PRIMA, Pixium Vision) have demonstrated the potential of photovoltaic subretinal prothesis to provide central vision in AMD patients with acuity closely matching the 100 μm pixels size^8,9^. These implants exhibited good durability in the subretinal space, stability of the retina above the implant and sustained visual acuity for over 4 years. To reduce the pixel size and thus enable higher visual acuity, we have proposed the use of 3-D honeycombs with elevated return electrodes, where penetration depth of electric field is defined by the wall height rather than pixel width^10^. Our earlier studies demonstrated migration of the inner retinal neurons into the honeycomb wells and their survival has been shown up to 6 weeks post implantation^13^. However, it is important to characterize the long-term structural and functional effects of such 3-D implants on the retina before translating this technology into clinical testing.

RCS rats used in this study constitute a good model for studying retinal structure and response to electrical stimulation after photoreceptor degeneration. The age-related retinal thinning described in this study is also common to normally-sighted rats (decrease by up to 29%) ^19^ and humans (by up to 39%)^20–22^. Data with AMD patients suggest that after initial thinning during the first 3 months post-op, retinal thickness changed very little during the 4 years follow-up ^23^. More significant thinning in RCS rats could be due to the different form of retinal degeneration, modeling RP rather than AMD. Importantly, there was no significant change in the retinal thickness between the clinically tested planar configuration and the proposed 3-D honeycomb implants.

Interestingly, the VEP peak amplitude recorded on the day of implantation drops nearly 3-fold within a week, and then gradually recovers and eventually exceeds the day 0 measurements. Since proximity of the retina to the implant only improves with time, the signal decline during the first week and its gradual recovery cannot be explained by the improving proximity to the implant over time, especially since electric field generated by monopolar implants with full-field illumination penetrates deep into the retina, hence negating the distance issues. We hypothesize that the drop in VEP could be related to acute glial reactivity. The surgical insult, involving a scleral, choroidal and retinal incisions and the introduction of a foreign object in the subretinal space, results in microglia and systemic immune cell infiltration, migration and accumulation. Müller glia become reactive, increasing the GFAP expression, cellular swelling, migration of nuclei and potentially forming a glial seal. The acute glial response sets in within hours, reaching its peak within the first week. As the acute reactivity slowly subsides and the retina integrates with the implant, and the VEP signal starts to recover. However, this drop is unlikely to affect patients since they start visual rehabilitation and training after about 3 months post-op.

Stimulation threshold was not affected by 40 μm honeycombs, compared to planar implants, but with 20 μm honeycomb pixels, it significantly increased after 18 weeks post-op. Since the number of cells in the INL with 20 μm pixels did not differ much from that with 40 μm pixels, the difference may be related to cellular excitability. It is possible that diffusion of oxygen and nutrients within tighter wells is more restricted, resulting in a decrease of the bipolar cell function. Another plausible explanation could be related to reduced penetration of Müller cells into 20 μm compared to 40 μm wells, visualized by their SOX9 positive nuclei, glutamine synthetase (GS) and GFAP staining^13^. Bipolar cells rely more heavily on Müller cells for their homeostatic and metabolic support than other retinal neurons^24^, and hence decreased interaction between bipolar and Müller cells inside tighter wells results in their decreased excitability.

One way to improve the oxygenation and allow better Müller cell migration into the wells is to decrease the wall height from 25 μm to about 15 μm. However, lower walls would decrease the electric penetration into the retina. The 15 μm deep wells should still accommodate two layers of bipolar cell nuclei, representing about half of the INL thickness, which is sufficient for eliciting retinal response^10^, but the percepts might not be as bright as with the full-depth stimulation.

Stimulation thresholds of 0.05 mW/mm^2^ with 40 μm 0.08 mW/mm^2^ with 20 μm honeycomb pixels are significantly lower than 1.8 mW/mm^2^ with 40 μm flat bipolar pixels^12^, confirming the promise of the honeycomb-based approach to high resolution subretinal prosthesis. The 40 μm pixels could theoretically provide visual acuity of 20/160, which exceeds the US legal blindness limit of 20/200, and with 20 μm pixels it may exceed 20/100. However, since AMD patients have a thicker ‘debris’ layer between the INL and the implant ^8,23^ than in RCS rats, further optimization will be required before clinical tests of this technology. Developing an animal model of AMD, which includes this debris layer, will be essential for assessment of migration the bipolar cells into the wells.

## Methods

### Fabrication of 3-dimensional implants

To fabricate the 3D honeycombs on flat arrays of photodiodes, 4 μm wide and 25 μm tall walls on the circumference of each pixel were built by two-photon polymerization using Nanoscribe Photonics GT (Nanoscribe GmbH & Co., Eggenstein, Germany). The dip-in laser lithography was performed using a 25x objective (NA=1.4) in IP-S photoresist. The layouts of the flat monopolar implants with 40 μm and 20 μm pixels^25^ were constructed in SolidWorks (Dassualt Systems, France) and a hexagonal mesh of walls was extruded from the metal traces between the pixels, corresponding to return electrodes, using the STL protocol (Figure 1). To avoid overheating and improve adherence to the metal traces, laser power was set to 30% and a Galvo scanning rate to 5 mm/s for the first 5 μm height of the polymer walls, whereas the rest was polymerized at 40% power and 16 mm/s scanning rate.

### Surgical procedure and animal handling

All experimental procedures described in this work were conducted in accordance with the Statement for the Use of Animals in Ophthalmic and Vision research of the Association for Research in Vision and Ophthalmology (ARVO) and approved by the Stanford Administrative Panel on Laboratory Animal Care (APLAC protocol 13765 and 33394). Royal College of Surgeons (RCS) rats were used as an animal model of inherited retinal degeneration. Animals were maintained at the Stanford Animal Facility under 12h light/12h dark cycles with food and water ad libitum.

Animals were anesthetized with a mixture of ketamine (75 mg/kg) and xylazine (5 mg/kg) injected intraperitoneally. The photovoltaic devices were implanted in the subretinal space typically at 6 months of age, after a complete loss of the outer nuclear layer, as evidenced by optical coherence tomography (OCT; HRA2-Spectralis; Heidelberg Engineering, Heidelberg, Germany). Further surgical details are described in our previous work^26^. The implants were placed in the temporal-dorsal region, approximately 1 mm away from the optic nerve. A total of 17 animals were implanted with 1.5 mm diameter arrays: 3D implants with pixel sizes of 40μm (n=6) and 20μm (n=5), as well as flat implants of the same pixel sizes (n=6), as illustrated in Figure 1. Since there are almost no structural differences and no stimulation threshold difference between flat implants of different pixel sizes, flat arrays with both pixel sizes are grouped together as a single control group.

To visualize the retina and the implant, animals were monitored over time via OCT. The retina above the wells were measured using the Heidelberg software thickness analysis command. For electrophysiological measurements of the visually evoked potentials (VEPs), each animal was implanted with three transcranial screw electrodes: one electrode over each hemisphere of the visual cortex (4mm lateral from midline, 6mm caudal to bregma), and a reference electrode (2mm right of midline and 2mm anterior to bregma).

### Electrophysiological measurements

For each electrophysiological measurement session, the rat was anesthetized, pupil of the implanted eye was dilated, and its cornea was covered with a viscoelastic gel and a cover slip, to cancel its optical power and ensure good retinal visibility. Subretinal implant was illuminated with a customized projection system, consisting of an 880nm laser (MF-880nm-400μm, DILAS, Tucson, AZ), customized optics, and a digital micromirror display (DMD; DLP Light Commander; LOGIC PD, Carlsbad, CA). The entire optical system was integrated with a slit lamp (Zeiss SL-120; Carl Zeiss, Thornwood, NY) for real-time observation of the illuminated retina via a NIR-sensitive CCD camera (acA1300-60gmNIR; Basler, Ahrensburg, Germany). Light intensity at the cornea was calibrated during each session, and then scaled to the retinal plane by the ratio of a projected square on the retina and in air, to derive the NIR irradiance on the implant.

VEPs were recorded across the ipsilateral and contralateral visual cortices via the Espion E3 system (Diagnosys LLC, Lowell, MA) at a sampling rate of 2kHz, averaged over 250 trials. Corneal signal was simultaneously measured using ERG electrodes on the cornea and a reference electrode in the nose, while the ground electrode was placed in the rat’s tail. Corneal signal also served as a template for a stimulus artifact removal in the VEP waveforms. We applied the same stimulation protocols to all 3 groups of implants: 3D devices with 40μm and 20μm pixels, as well as flat devices.

For stimulation threshold measurements, full-field illumination was projected at 2 Hz with 10ms pulses, at varying irradiance levels, ranging from 0.002 to 4.7 mW/mm^2^ on the implants. The noise floor was determined by measuring VEP without any illumination. Threshold was defined as the first sampling point where the peak-to-peak VEP amplitude rises above the noise floor without any interpolation, as described previously^11^.

### Whole-mount retinal immunohistochemistry

For the retinal imaging and analysis, animals were euthanized 9-12 months post implantation using an intracardiac injection of Phenytoin/pentobarbital (Euthanasia Solution; VetOne, Boise, ID, USA). The eyes were enucleated and rinsed in phosphate buffered saline (PBS), anterior segment and lens were removed, the eye cup was cut to a 3 × 3 mm square centered around the implant, and fixed in 4% paraformaldehyde (PFA; EMS, PA, USA) for 12 hours at 4° C. Samples were permeabilized with 1% Triton X-100 (Sigma-Aldrich, CA, USA) in PBS for 3 hours at room temperature, followed by a blocking step in 10% bovine serum albumin (BSA) for 1 hour at room temperature, and a 48-hour incubation at room temperature with primary antibodies (Supplementary table 1) in 0.5% Triton X-100, 5% BSA in PBS. Samples were washed for 6 hours at room temperature in 0.1% Tween-20 in PBS (PBS-T), incubated for 48 hours at room temperature with secondary antibodies (Supplementary table.1) and counterstained with 4’, 6-Diamidin-2-phenylindol (DAPI) in PBS. After 6 hours of washing in PBS-T, the samples were mounted with Vectashield medium (H-1000; Vector Laboratories, Burlingame, CA, USA).

3-D imaging of the retinal whole mounts was performed using a Zeiss LSM 880 Confocal Inverted Microscope with Zeiss ZEN Black software. The implant surfaces were identified by reflection of a 514 nm laser with a neutral-density beam splitter allowing 80% transmission and 20% reflection. The images were acquired through the total thickness of the retina using a Z-stack, with the upper and lower bounds defined at the inner limiting membrane (ILM) and 10 μm below the base of the honeycomb wells, respectively. Stacks were acquired in the center of each honeycomb quadrant using a 40x oil-immersion objective with acquisition area >225×225 μm and 380 - 470 nm z-steps. The Zeiss z-stack correction module was used to account for dimmer light within the wells of the implants.

Confocal fluorescence datasets were processed using the FiJi distribution of ImageJ^27^. To correct for brightness variations at different Z positions in the stack within the wells and above the implant, we first maximized the contrast in the individual XY planes to ensure 0.3% channel saturation. The XY planes were de-speckled with the median filter and the background was suppressed using the rolling-ball algorithm^28^. The images then underwent cascades of gamma adjustments and min-max corrections to further suppress the background, depending on the noise level. Gaussian blurring was applied for nucleus staining channels to smoothen the brightness variations within individual cells. The implants were reconstructed by extruding the implant reflection toward the bottom (extraocular side) of the image stack.

To quantify the number of cells in the wells, the total DAPI and the DAPI within the wells were were was segmented into voxels based on the reflection channel using the Moore-Neighbor tracing algorithm implemented by the “bwboundaries” function in MATLAB 2021b (Mathworks, Inc., Natick, MA), while a control image stack (without an implant) was treated as one segment. Voxels brighter than 15% of the maximum intensity were considered positive, and in each segment, we defined the filling percentage as a fraction of positive voxels.

## Acknowledgements

We would like to thank Dr. Tiffany Huang for her contribution to implant fabrication. MB performed surgeries, in-vivo imaging, immunohistochemistry and confocal imaging; BW, MB conducted the electrophysiological measurements; MB, DPH, SS conducted image processing and image quantification; AS, LG and CZC fabricated subretinal implants under the guidance of KM and TK; DP guided the research and data analysis; all authors participated in writing and/or reviewing the manuscript. Studies were supported by the National Institutes of Health (Grants R01-EY-027786, and P30-EY-026877), the Department of Defense (Grant W81XWH-19-1-0738), AFOSR (Grant FA9550-19-1-0402), International Retinal Research Foundation, Wu Tsai Institute of Neurosciences at Stanford, and unrestricted grant from the Research to Prevent Blindness. Photovoltaic arrays were fabricated at the Stanford Nano Shared Facilities (SNSF) and Stanford Nanofabrication Facility (SNF), which are supported by the National Science Foundation award ECCS1542152. K.M. was supported by a Royal Academy of Engineering Chair in Emerging Technology, UK.

## Notes

### Competing Interest Statement

Daniel Palanker and Ted Kamins- consultant (Pixium Vision)
Daniel Palanker - Patent (Stanford University and Pixium Vision)

