## Supplementary figures and images for "3D electronic implants in subretinal space: long-term follow-up in rodents"

### Supplementary table 1

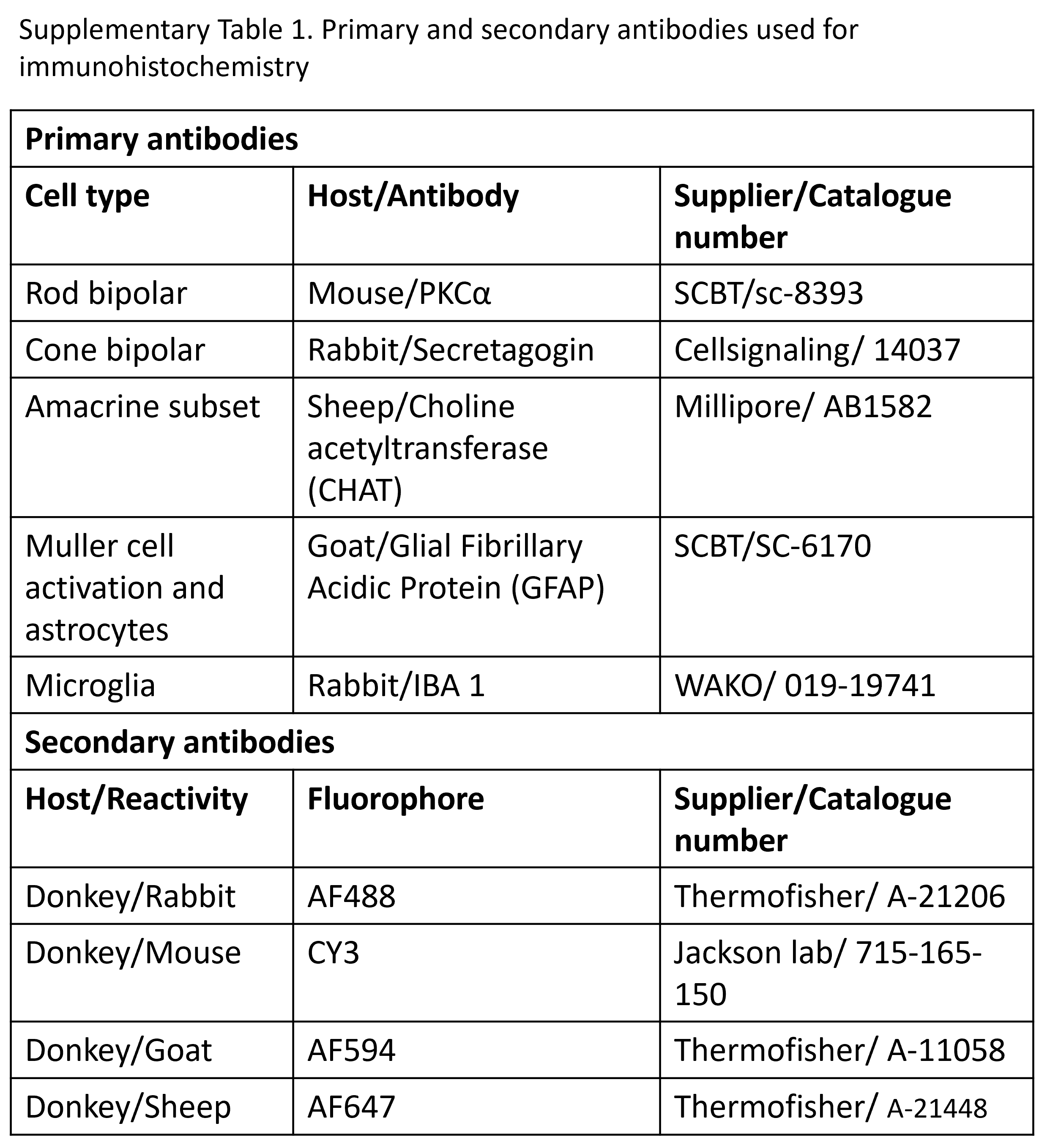
